# Multi-modal transcriptomic analysis unravels enrichment of hybrid epithelial/mesenchymal state and enhanced phenotypic heterogeneity in basal breast cancer

**DOI:** 10.1101/2023.09.30.558960

**Authors:** Sarthak Sahoo, Soundharya Ramu, Madhumathy G Nair, Maalavika Pillai, Beatriz P San Juan, Heloisa Zaccaron Milioli, Susmita Mandal, Chandrakala M Naidu, Apoorva D Mavatkar, Harini Subramaniam, Arpita G Neogi, Christine L Chaffer, Jyothi S Prabhu, Jason A Somarelli, Mohit Kumar Jolly

## Abstract

Intra-tumoral phenotypic heterogeneity promotes tumor relapse and therapeutic resistance and remains an unsolved clinical challenge. It manifests along multiple phenotypic axes and decoding the interconnections among these different axes is crucial to understand its molecular origins and to develop novel therapeutic strategies to control it. Here, we use multi-modal transcriptomic data analysis – bulk, single-cell and spatial transcriptomics – from breast cancer cell lines and primary tumor samples, to identify associations between epithelial-mesenchymal transition (EMT) and luminal-basal plasticity – two key processes that enable heterogeneity. We show that luminal breast cancer strongly associates with an epithelial cell state, but basal breast cancer is associated with hybrid epithelial/mesenchymal phenotype(s) and higher phenotypic heterogeneity. These patterns were inherent in methylation profiles, suggesting an epigenetic crosstalk between EMT and lineage plasticity in breast cancer. Mathematical modelling of core underlying gene regulatory networks representative of the crosstalk between the luminal-basal and epithelial-mesenchymal axes recapitulate and thus elucidate mechanistic underpinnings of the observed associations from transcriptomic data. Our systems-based approach integrating multi-modal data analysis with mechanism-based modeling offers a predictive framework to characterize intra-tumor heterogeneity and to identify possible interventions to restrict it.

## Introduction

Intra-tumoral heterogeneity in breast cancer remains a key obstacle in the effective management of the disease (Marusyk et al., 2020). A major determinant of molecular heterogeneity in breast cancer is attributed to molecular subtype characteristics, which can be broadly classified as luminal or basal (Skibinski and Kuperwasser, 2015). In addition, cancer cells can exhibit different interconvertible cellular states along varied axes of plasticity such as epithelial-mesenchymal transition (EMT), stemness, metabolic reprogramming, and immune evasion traits to create an increased overall phenotypic heterogeneity (Brown et al., 2022; Hong et al., 2018; Jain et al., 2022; Wahl and Spike, 2017). The extent of the crosstalk among these different axes, which is often mediated via feedback loops, can have major implications in dependence and coordination between plasticity axes on each other as well as overall disease progression (Jia et al., 2021; Sahoo et al., 2021a). Specifically, in breast cancer, two mainstays of molecular heterogeneity that are often used interchangeably are the luminal-basal and epithelial-hybrid-mesenchymal states (Lesniak et al., 2013). This assumed equivalence, based at least partly on gene set enrichment analysis, largely considers EMT as a binary process (Prat et al., 2010). However, it has now been extensively reported that EMT in breast cancer exists more as a spectrum of phenotypes residing along the epithelial-mesenchymal axis (Bierie et al., 2017; Grosse-Wilde et al., 2015; Jolly et al., 2017; Yu et al., 2013). Therefore, the association of luminal-basal lineage characteristics and associated plasticity with a partial EMT (pEMT) or hybrid epithelial/mesenchymal (E/M) plasticity remains largely unclear (Mohamed et al., 2023). Similarly, the extent to which associations between partial/full EMT and luminal-basal plasticity in breast cancer are generalizable also remain to be elucidated.

Current therapeutic approaches often target specific molecular subtypes of breast cancer (Ignatiadis and Sotiriou, 2013; Yin et al., 2020). However, therapy-driven adaptive plasticity and consequent phenotypic heterogeneity pose challenges in achieving durable responses (Qin et al., 2020). In addition, the clinical implications of understanding coupling between EMT and luminal-basal plasticity, and its impact on the estrogen receptor (ER) signalling, in breast cancer are substantial. Intra-tumor heterogeneity and phenotypic plasticity can impact treatment response and patient survival by altering the dependence of breast cancer cells on a luminal program and ER signalling (Lindström et al., 2018; Mohamed et al., 2023; Sahoo et al., 2021a; Yu et al., 2021). Heterogeneity can also drive collective behaviour by facilitating division of labour among different cancer cell subpopulations, and consequently varying interactions with the surrounding microenvironment (Bhattacharya et al., 2021). Thus, higher phenotypic heterogeneity can have both cell-autonomous and non-cell-autonomous contributions to population-level fitness. Gaining insights into the underlying dynamics and mechanisms of breast cancer cell heterogeneity is critical to understanding plasticity-associated therapy relapse (Burkhardt et al., 2022; Pillai et al., 2023).

Here, we uncover the associations between epithelial-hybrid-mesenchymal cell-state and the luminal-basal axis using multi-modal transcriptomics (bulk, single cell and spatial transcriptomics) data from breast cancer cell lines and primary tumor samples. Our analysis demonstrates that luminal cell lines and tumors strongly associate with an epithelial phenotype, but basal cell lines and tumors are not fully mesenchymal. Instead, basal breast cancer associates with pEMT phenotype as well as an enriched phenotypic heterogeneity along the EMT spectrum. These patterns were also recapitulated in methylation profiles, indicating an epigenetic crosslinking between EMT and lineage plasticity along the luminal-basal axis. Finally, we propose an underlying gene regulatory network including players mediating EMT and luminal and basal differentiation axes. The emergent dynamics of this network could recapitulate the trends observed in transcriptomic data, offering novel insights into molecular underpinnings of basal breast cancer. Our integrative systems biology approach reveals hallmarks of basal breast cancer heterogeneity (enrichment of hybrid E/M phenotypes, and higher diversity in terms of EMT cell-states) and proposes a mechanistic computational model of crosstalk between EMT and lineage plasticity as a framework to test possible therapeutic interventions to restrict intra-tumor heterogeneity.

## Results

### A luminal gene expression signature is closely associated with an epithelial state

Clinical and molecular classification of breast cancer has yielded key insights into breast cancer disease biology and has provided a much-needed stratification of the disease for its effective management in a clinical setting (Łukasiewicz et al., 2021). However, the activities of the various biological pathways/processes, including the level of activity of the various gene sets corresponding to the breast cancer subtypes, contributing to the overall phenotypes exhibited by cancer cells can still be quite varied even within a well stratified breast cancer subtype (Turner et al., 2021). While such heterogeneities can significantly impact the disease outcome, the origins and molecular underpinnings of said heterogeneities remain largely unexplored. Hence, we sought to uncover associations relating to two key biological axes of plasticity – the extent to which breast cancer samples express the luminal/basal gene expression programs and the extent to which the samples are epithelial/mesenchymal, whether we examine a particular subtype of breast cancer or consider the entire cohort of subtypes as a unified group.

To uncover the associations between the luminal-basal phenotypes and the EMT status of breast cancer cells, we sought to compare how these four gene sets - luminal, basal, epithelial and mesenchymal - correlated with one another independently. To do this, we first calculated single sample gene set enrichment analysis (ssGSEA) scores for corresponding gene expression signatures to quantify the activity of these four gene sets. The epithelial and mesenchymal gene signatures were adapted from a previous pan-cancer analysis (Tan et al., 2014). The luminal and basal nature of breast cancer samples was assessed using a previously curated set of 15 luminal and basal specific genes each (Nair et al., 2022). There was minimal overlap between the genes included in these signatures (**Table S1**). As expected, in both CCLE cohort of cell lines and in TCGA breast cancer samples, which cover all major breast cancer subtypes (luminal A, luminal B, HER2+ and basal), the epithelial signature score correlated negatively with the mesenchymal score (CCLE: r = - 0.73, p < 0.01, TCGA: r = -0.414, p < 0.01), and the luminal signature score correlated negatively with the basal score (CCLE: r = -0.331, p < 0.05, TCGA: r =-0.369, p < 0.05) (**Fig S1A-B**).

In CCLE, we observed a positive correlation of epithelial signature with the luminal signature (r = 0.657, p < 0.01) (**Fig 1A, i**). Conversely, the luminal signature correlated negatively with the mesenchymal score (r = -0.775, p < 0.01) (**Fig S1C, i**), indicating that the luminal nature was strongly associated with an elevated epithelial and reduced mesenchymal state. The basal signature, on the other hand, showed relatively weaker trends. It correlated positively with the mesenchymal nature (r = 0.409, p < 0.01) (**Fig 1A, ii**), but showed no significant correlation with the epithelial nature of cell lines (**Fig S1C, ii**). These observations corroborate previous reports of basal cells expressing some mesenchymal markers (VIM, SLUG) (Gudjonsson et al., 2005; Phillips and Kuperwasser, 2014), while also considered as a part of mammary epithelium in normal tissues and in breast cancer (Gusterson et al., 2005; Nguyen et al., 2018).

**Figure 1:**
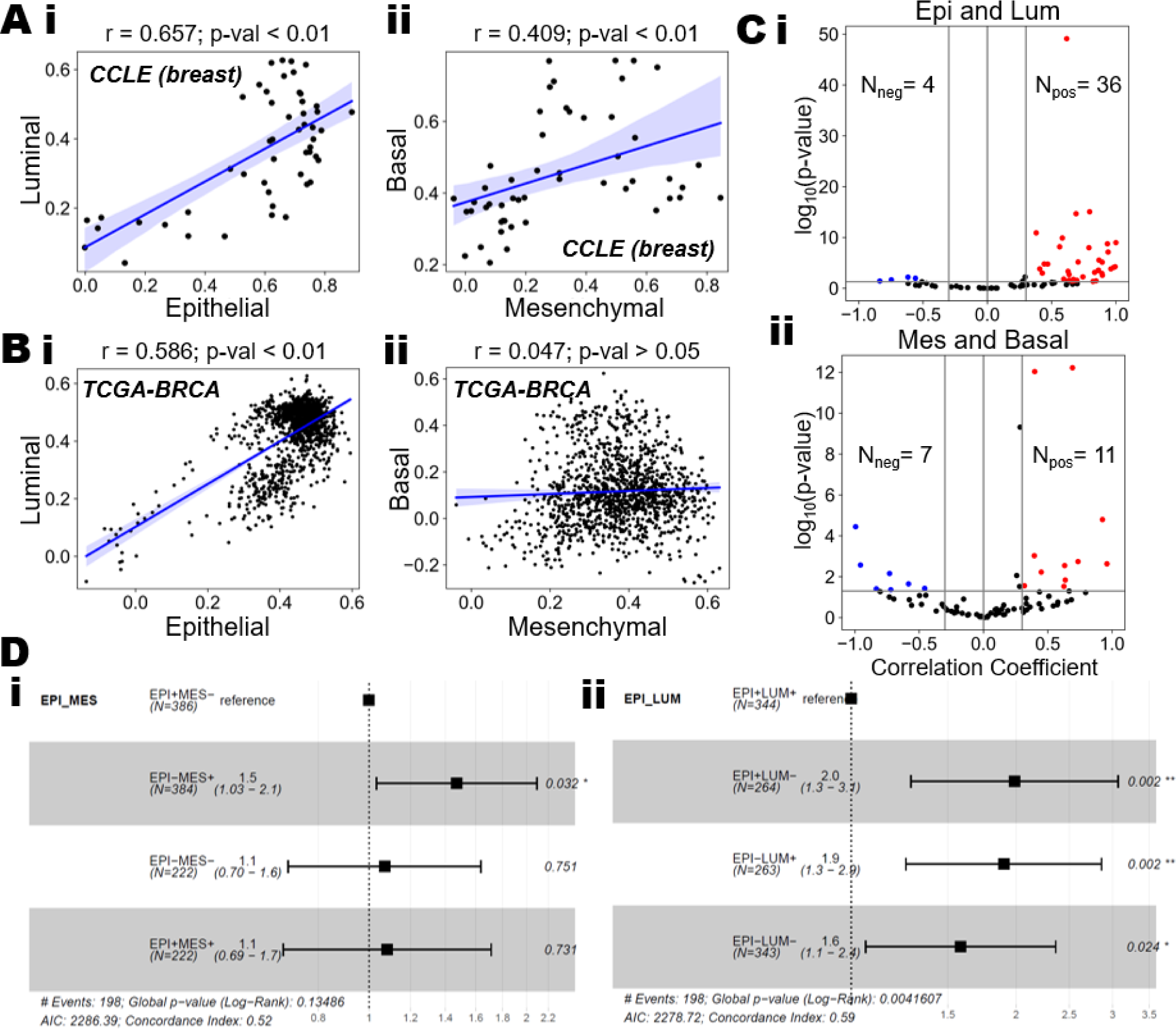
Associations between luminal-epithelial and basal-mesenchymal programs in breast cancer. **A)** Scatterplots showing the correlations between ssGSEA scores of **i)** Epithelial-Luminal and **ii)** Basal-Mesenchymal programs in CCLE breast cancer cell lines. Pearson’s correlation coefficients are reported with the corresponding p-values. **B)** Same as A) but for TCGA BRCA patients. **C)** Meta analysis of 80 breast cancer specific datasets showing volcano plots of correlation coefficient values for **i)** Epithelial-Luminal and **ii)** Basal-Mesenchymal programs. Pearson’s correlation coefficients and the corresponding p-values are shown. **D)** Forest plots showing survival analysis in TCGA breast cancer patients stratified based on their ssGSEA scores on the **(i)** epithelial-mesenchymal spectrum and **(ii)** epithelial-luminal spectrum. The reference cell population is set to high epithelial, low mesenchymal (EPI+ MES-) and high epithelial, high luminal (EPI+, LUM+) states, respectively. * represents a 0.01 < p-value < 0.05 while ** represents a p-value < 0.01.

Next, we examined if the above trends were observed in primary tumor samples. Thus, we analysed the TCGA breast cancer cohort, considering all subtypes – Luminal A, Luminal B, HER2+ and Basal-like. Similar to cell line data, the luminal signature positively correlated with the epithelial signature (r = 0.657, p < 0.01) (**Fig 1B, i**). However, in contrast to the cell line data, the basal nature was not correlated with the mesenchymal phenotype (r = 0.047, p > 0.05) (**Fig 1B, ii**), but showed negative correlation trends with the epithelial nature (r=-0.259, p < 0.01) (**Fig S1D, i**). Further, the luminal nature was not significantly correlated with the mesenchymal signature (**Fig S1D, ii**). Upon quantifying the different correlations especially in the Luminal A and the Basal-like subtypes, we found that the luminal-epithelial association was consistently positive and significant in both cases (**Table S2**). Further, in the Luminal A subtype patients, the following was observed: a) mesenchymal score does not correlate with the basal score (r = 0.16, p < 0.01) or with the luminal score (r = -0.19, p < 0.01), and b) the basal score correlates positively with epithelial score (r = 0.35, p < 0.01), suggesting that within luminal A subtype, tumors enriched for a basal character were likely to be both high in epithelial and mesenchymal nature. Similar trends, albeit to a weaker extent, were noted in the basal-like subtype (**Table S2**).

Having shown that the luminal-epithelial association is stronger than a basal-mesenchymal association in a representative cell line dataset (CCLE) and patient samples (TCGA), we examined whether these trends were more broadly conserved in other breast cancer datasets. We performed a meta-analysis of the associations between these pairs of pathways in 80 transcriptomic datasets (**Table S3**). We observed that in 36 datasets the correlations between luminal and epithelial signatures were significant and positive (r > 0.3, p < 0.05) while only 4 datasets showed a significant negative trend (r < -0.3, p < 0.05) (**Fig 1C, i**). Such a skew towards the positive side was not present for the correlations between the basal and the mesenchymal signatures (11 vs 4 respectively) (**Fig 1C, ii**). Similarly, such large skews were not found for the luminal-mesenchymal pair or the epithelial-basal pair (**Fig S1E**). These results indicate that the association between the luminal and epithelial signatures is more common across breast cancer samples both *in vitro* and *in vivo* as compared to those between the basal and mesenchymal signatures.

To determine if the association between the epithelial and luminal programs can provide additional power to stratify breast cancer patients, we compared the prognostic capacity of an epithelial-mesenchymal gene set with the epithelial-luminal gene set in the TCGA breast cancer cohort. Using the epithelial-mesenchymal signatures we observed that the patients with high mesenchymal (EPI-MES+) tumors showed a poorer prognosis (Hazard ratio = 1.5, p < 0.05) (**Fig 1D, i**). The groups of patients with tumors belonging to mixed epithelial and mesenchymal characteristics (EPI+MES+ and EPI-MES-) were not well segregated from the reference group. However, when we performed the analysis with our epithelial-luminal classification, we found that with respect to the reference distribution of patients i.e., patients with high epithelial and high luminal tumors (EPI+LUM+), all the other groups showed significant differences in survival (**Fig 1D, ii**). Patients with tumors showing low luminal and/or epithelial signatures (EPI+LUM-, EPI-LUM+, EPI-LUM-) had worse prognosis. Collectively, these results indicate a positive correlation between the luminal and epithelial programs that can be used effectively to stratify patients into prognostic survival groups.

### Basal-like breast cancer samples are associated with a partial EMT signature in bulk transcriptomic datasets

Given the unclear nature of association of the basal nature with either the epithelial or mesenchymal markers signatures exclusively, we postulated that the basal signature may be correlated with a pEMT signature. We used the pEMT signature reported earlier (Puram et al., 2017) to estimate the pEMT nature of the bulk samples by calculating the ssGSEA scores.

In the CCLE breast cancer cell line set, we observed that the pEMT signature was positively correlated with basal signature (r = 0.603, p < 0.01) (**Fig 2A**) and, to a lesser extent, with the basal-mesenchymal correlation previously observed (r = 0.409, p < 0.01) (**Fig 1A, ii**). Similarly, in TCGA samples, the correlation of pEMT and basal signatures was higher than that of a mesenchymal signature with basal (r = 0.153, p < 0.01 vs. r = 0.047, p > 0.05 respectively) (**Fig 2B** and **Fig 1B, ii**). We next investigated these trends in a subtype-specific manner and found a numerically higher association of the basal signature with a pEMT state rather than a mesenchymal state (compare **Fig 2B, ii** with **Fig 2B, i;** compare **Fig 2C, ii** with **Fig 2C, i**). In other words, within the luminal A and B subtypes, as the basal nature increases, a consistent upregulation of a pEMT state, but not necessarily that of mesenchymal state, is observed.

**Figure 2:**
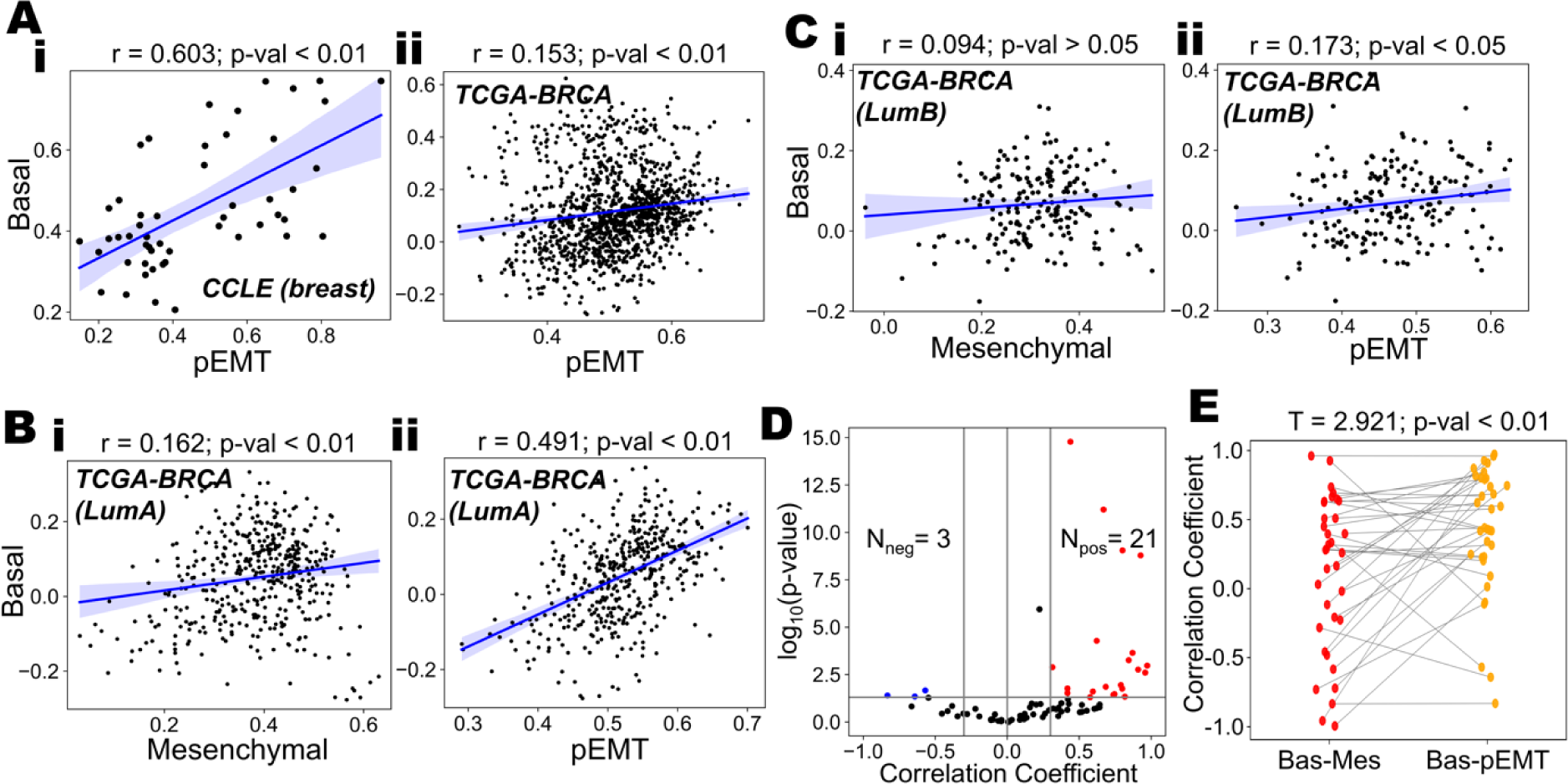
Associations between pEMT and basal programs in breast cancer. **A)** Scatterplots showing the correlations between the ssGSEA scores of pEMT and Basal in **(i)** CCLE breast cancer cell lines and **(ii)** TCGA breast cancer patients. Pearson’s correlation coefficients have been reported with the corresponding p-values. **B)** Scatterplots showing the correlations between the ssGSEA scores of **(i)** Mesenchymal-Basal and **(ii)** pEMT-basal programs in Luminal A subtype of TCGA BRCA patients. Pearson’s correlation coefficients have been reported with the corresponding p-values. **C)** Scatterplots showing the correlations between the ssGSEA scores of **(i)** Mesenchymal-Basal and **(ii)** pEMT-basal programs in Luminal B subtype of TCGA BRCA patients. Pearson’s correlation coefficients have been reported with the corresponding p-values. **D)** Meta analysis of 80 breast cancer specific datasets showing volcano plots of correlations for pEMT and basal programs. Pearson’s correlation coefficients and the corresponding p-values have been used. **E)** Pair plot showing the comparison of the Pearson correlation coefficients between the Basal-Mesenchymal and the Basal-pEMT programs. Paired T test has been performed and the T statistic and p-value have been reported.

Next, we interrogated whether the association of the pEMT and basal signatures was observed more broadly in breast cancer samples. In our meta-analysis of 80 datasets (across many breast cancer subtypes), we observed that the basal signature was positively correlated with the pEMT state in 21 of them (r > 0.3, p < 0.05) but negatively in only 3 of them (r < -0.3, p < 0.05) (**Fig 2D**). This skew towards a positive correlation is greater than the one observed for the basal-mesenchymal pair (**Fig 1C, ii**). These results indicate that irrespective of the clinical/molecular subtype of breast cancer samples/cell lines, there exists a positive association between the basal nature and the pEMT nature of samples. We also compared the Pearson’s correlation coefficients of basal-pEMT set with those of basal-mesenchymal set in a paired manner and found a significant increase in basal-pEMT correlation (T = 2.9, p < 0.01) (**Fig 2E**). These results collectively suggest that at a bulk transcriptomic level, breast cancer samples show a stronger association of basal-like breast cancer with a pEMT signature instead of a mesenchymal one.

These trends are further supported by recent experimental observations that luminal progenitors, the proposed cell of origin of basal-like tumors, undergo a pEMT at onset of tumorigenesis (Landragin et al., 2022). Moreover, mammary basal epithelial cells have been shown to exhibit a pEMT state, i.e. co-expressing typical epithelial and mesenchymal markers – *ZEB1* and *OVOL2* respectively (Han et al., 2022). Furthermore, genetically engineered mouse models, patient-derived xenografts and patient samples of triple negative breast cancer cells demonstrated large populations of hybrid E/M cells *in vivo* that lead invasion (Grasset et al., 2022). Together, these observations underscore the association of the basal breast cancer cell state with a pEMT phenotype.

### Epigenetic status of epithelial and mesenchymal genes underlies associations between the luminal-epithelial and basal-pEMT programs

Having shown the close association of a luminal program with an epithelial state, and that of basal program with a pEMT state at a bulk transcriptomic level, we wished to interrogate whether these associations may have an epigenetic basis. Thus, we analysed genome-wide methylation data of CCLE breast cancer cell lines (GSE42944) and compared it with the corresponding gene expression data. We observed that the cell lines that were primarily classified into three groups – Luminal, Basal A, and Basal B (Cope et al., 2014) – were distinctly positioned on the two-dimensional EMT plane (**Fig 3A, i**). The luminal subtype of cell lines clustered towards the (high epithelial, low mesenchymal) part of the plane (**Fig 3A, i**), suggesting a strong association with the epithelial characteristics. Conversely, the Basal B cell lines positioned diametrically opposite, aligning with the mesenchymal end. The Basal A subtype, however, occupied intermediate regions of the EMT plane, indicating a mixed epithelial-mesenchymal profile. Consistent with our previous observations, the basal score of the cell lines was negatively associated (r = -0.34, p < 0.05) with their luminal score (**Fig 3A, ii**) while being positively associated with the pEMT score (**Fig S2A**).

**Figure 3:**
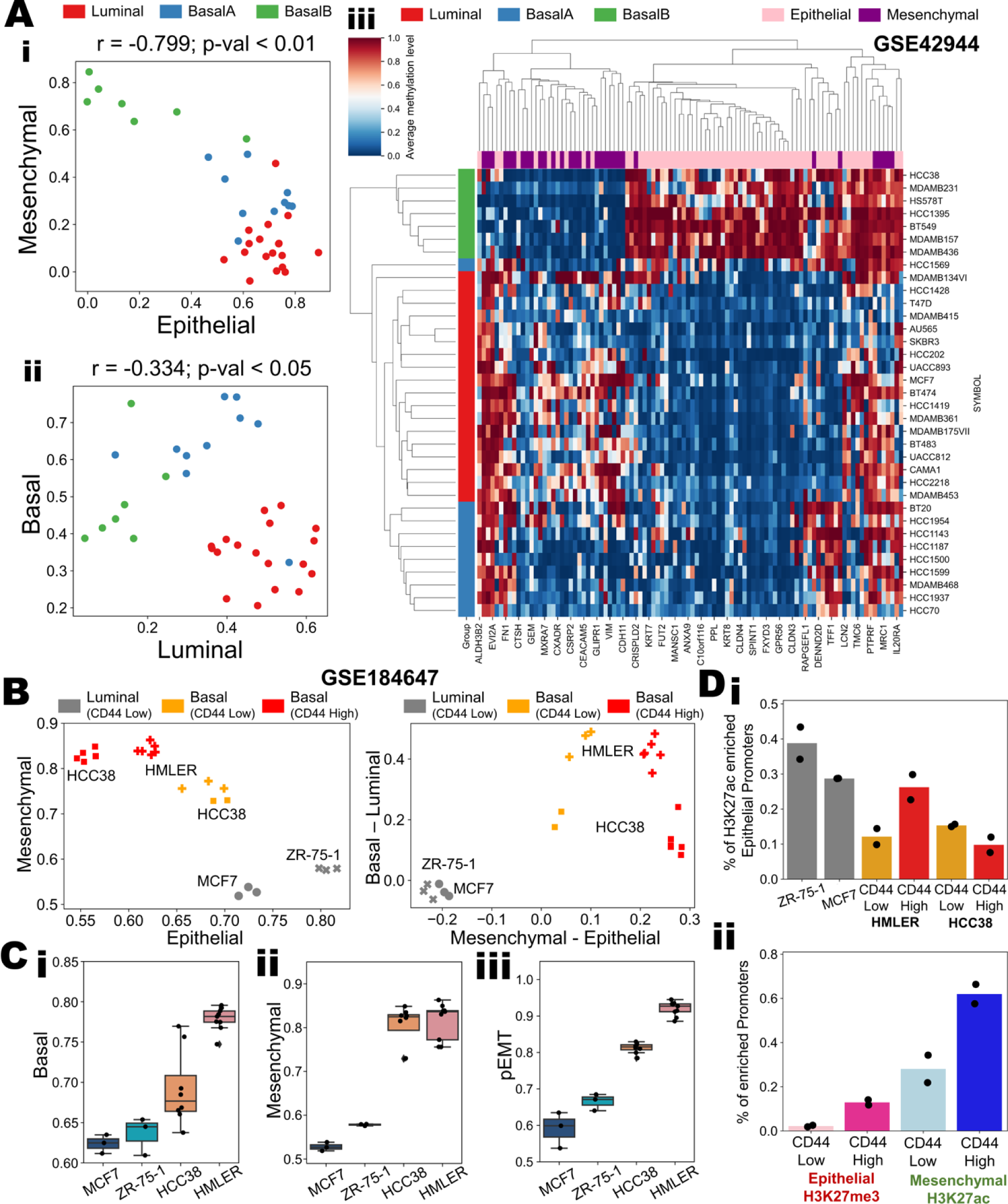
Epigenetic underpinnings of epithelial-mesenchymal phenotypes in luminal and basal breast cancer cell lines. **A)** Scatterplots for ssGSEA scores of **i)** Epithelial-Mesenchymal and **ii)** Luminal-Basal programs in breast cancer cell lines. Pearson’s correlation coefficient values and corresponding p-values are mentioned. **iii)** Heatmap showing the extent of methylation of CpG islands in the promoters of epithelial (pink) and mesenchymal (purple) genes for different breast cancer cell lines belonging to luminal (red), basal A (blue) and basal B (green) subtypes. **B)** Scatterplots showing the position of luminal, basal (CD44 low) and basal (CD44 high) cell lines on the two dimensional (left) epithelial-mesenchymal and (right) luminal-basal plane. **C)** Boxplots showing expression of **i)** Basal, **ii)** Mesenchymal and **iii)** pEMT gene expression programs across four different cell lines. **D) i)** Bar plots showing the percentage of H3K27ac activation marks in different breast cancer cell lines. **ii)** Bar plots showing the percentage of enriched promoters of epithelial inactivation marks and the mesenchymal activation marks across the CD44 high and low subpopulations in basal breast cancer cell lines.

We next quantified the methylation level for each gene from the epithelial and the mesenchymal gene sets across all cell lines belonging to the three subtypes. We observed that the luminal cell lines were extensively methylated in promoters of mesenchymal genes and had lower methylation levels on epithelial genes (**Fig 3A, iii**). The converse was true for the Basal B subtype of breast cancer cells. The Basal A subtype of breast cancer cells which were intermediate in terms of their EMT status exhibited lower levels of methylation in both the epithelial and mesenchymal genes (**Fig 3A, iii**). This indicates that while the Basal B subtype of breast cancer cells had mostly silenced epithelial genes, the Basal A subtype of cells have both epithelial and mesenchymal genes active that may explain the pEMT like phenotype observed. Furthermore, the basal cell lines (Basal A and Basal B taken together) exhibited higher levels of methylation of the *ESR1* gene (encoding for Estrogen receptor (ER)) (**Fig S2B, i**), a key gene contributing to luminal behaviour (**Fig S2B, ii**) compared to the more luminal cell lines. On the contrary, basal cell lines had lower methylation levels in genes belonging to the pEMT signature (**Fig S2B, iii**) and the ones belonging to basal signature (**Fig S2B, iv**).

Previous analysis has demonstrated that an isogenic cell line can have different subpopulations in terms of its EMT nature such as EpCAM-hi and EpCAM-lo subpopulations in PMC-42LA cells, or cells with varying CD24 and/or CD44 levels in multiple breast cancer cell lines (Bhatia et al., 2019; Deshmukh et al., 2021; Mani et al., 2008). Thus, we focused on heterogeneity within a cell line in terms of their epithelial-mesenchymal nature and their luminal-basal characteristics. We analysed the RNA-seq data of four cell lines (2 luminal – MCF7 and ZR-75, 2 basal – HCC38 and HMLER) (GSE184647). The luminal cell lines were CD44-low; thus, no CD44-high subgroup was observed in them, but basal cell lines harboured distinct CD44-low and CD44-high subpopulations (San Juan et al., 2022). We observed that luminal cell lines were clustered on the high epithelial, low mesenchymal section of the two-dimensional EMT spectrum, while the CD44-high subpopulation of basal cell lines were clustered towards the low epithelial, high mesenchymal end of it (**Fig 3B, left**). Intriguingly, the CD44 low basal subpopulations were clustered in the medium epithelial, medium mesenchymal region, indicative of a pEMT state (**Fig 3B, left**). We also observed that on the combined EMT and luminal-basal plane, the luminal cell lines were clustered together in the high epithelial, high luminal region, while the basal cell lines showed a larger spread, with the CD44-high subpopulations being more mesenchymal, but not necessarily more basal than their CD44-low counterparts as assessed by ssGSEA scores of the corresponding gene signatures (**Fig 3B, right**). We observed that HMLER cells were more basal compared to HCC38, i.e., basal nature was better explained by the cell line rather than CD44 (**Fig 3B, right**).

We also noted that luminal cell lines had lower ssGSEA scores for basal, mesenchymal and pEMT gene signatures as compared to basal cell lines (**Fig 3C**). Intriguingly, both the basal cell lines had distinct ssGSEA scores of basal gene set activity, but comparable ssGSEA scores for mesenchymal signature. The pEMT signature was better able to capture the trends of the basal signature compared to that of the mesenchymal signature (**Fig 3C**). This trend strengthens our observations that the pEMT signature is a better predictor of basal nature compared to that of a mesenchymal signature. Finally, we probed the MINT-CHIP data for these samples and found that the promoters of epithelial genes were specifically higher in H3K27ac (activation) marks for luminal cell lines compared to basal cell lines (**Fig 3D, i**). Furthermore, in HMLER, the CD44-low sub-population had consistently lower levels of activation marks (H3K27Ac) on mesenchymal genes as well as lower levels of suppressive marks (H3K27me3) on the epithelial genes (**Fig 3D, ii**). This pattern offers a potential explanation for the association of basal subtype with a pEMT state. Recent data from mammary stem cell subpopulations revealed higher accessibility and enrichment of P63 DNA-binding motifs in basal cells, and that of ELF5 DNA-binding motif in luminal progenitors (Dravis et al., 2018). Given the established role of NP63 in driving a pEMT program (Dang et al., 2015; Jolly et al., 2017; Westcott et al., 2020) and that of ELF5 in inhibiting EMT (Chakrabarti et al., 2012; Wu et al., 2015; Yao et al., 2015), these observations together support the possibility of an epigenetic control in the association of basal breast cancer with hybrid E/M phenotype(s).

### Spatial transcriptomics reveals intra-patient variability in EMT phenotypes in basal subtypes of breast cancer

After demonstrating a higher heterogeneity of basal breast cancer cell lines along the EMT spectrum *in vitro*, we sought to investigate the same in breast cancer patients. Thus, we analysed publicly available spatial transcriptomics datasets of breast tissue sections (Wu et al., 2021) to infer the patterns of spatial heterogeneity in associated luminal-epithelial and basal-mesenchymal/pEMT status, using therapeutically relevant markers of gene expression. Firstly, we observed that amongst the n=6 patients for whom spatial transcriptomics data was available, the ER+ breast cancer patients had distinctively higher levels of luminal nature but downregulated basal scores (**Fig 4A**). On the contrary, the basal breast cancer patients, while showing reduced levels of luminal nature, were quite heterogenous for basal signature expression (**Fig 4A**). The overall correlation for the epithelial-mesenchymal score pair was strongly negative (r = -0.85, p < 0.01) – while the ER+ breast cancer patients clustered towards the high epithelial, low mesenchymal portion of the plane, TNBC patients were highly variable and scattered across the spectrum (**Fig S2C**).

**Figure 4:**
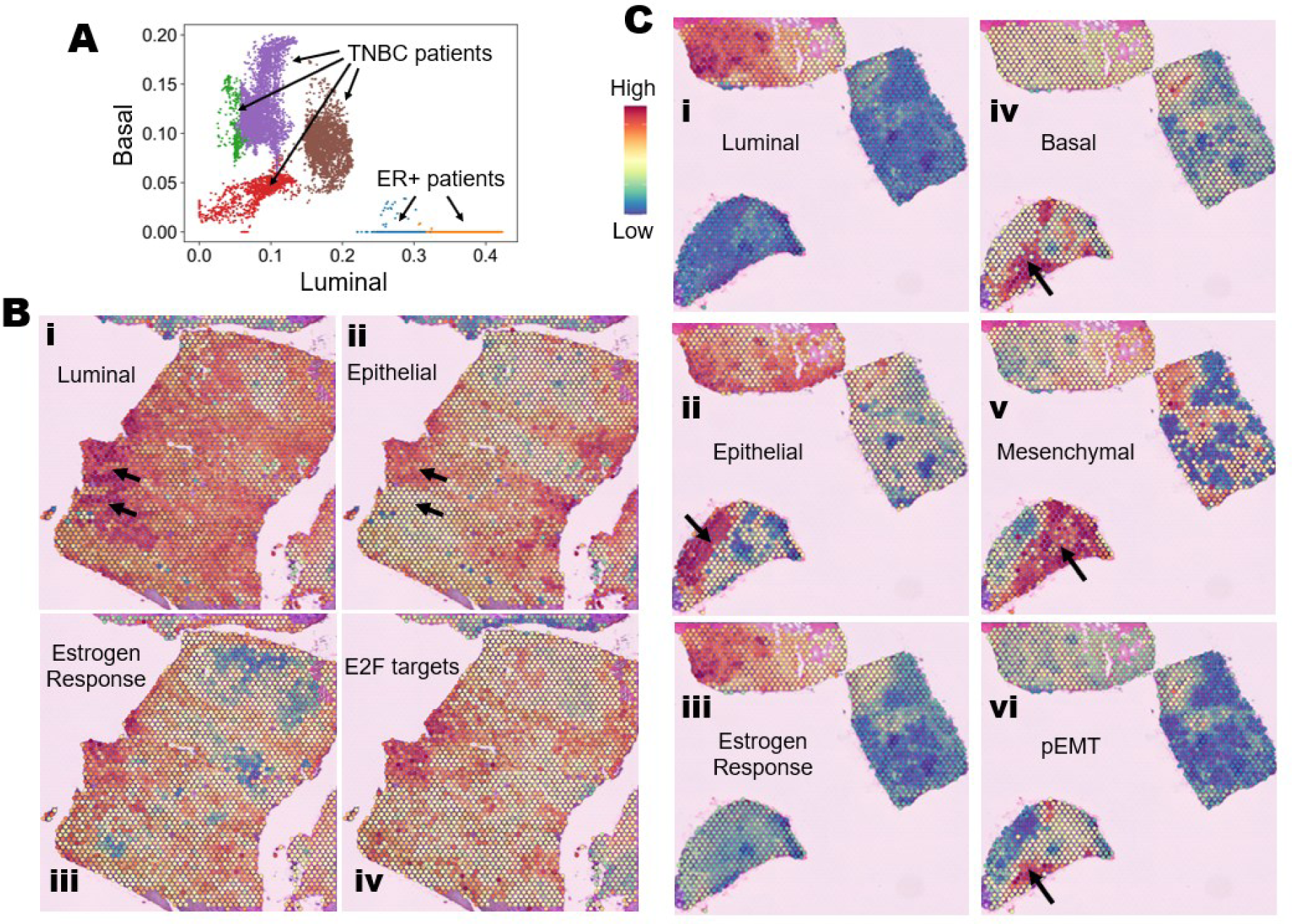
Spatial transcriptomic analysis of estrogen receptor positive and triple negative breast cancer patients. **A)** Scatterplot showing the position of the estrogen receptor positive (ER+) and triple negative breast cancer (TNBC) patients on a two dimensional luminal-basal plane. Each point the plane is based on the gene expression values for a specific spot on the spatial transcriptomic datasets. **B)** Spatial transcriptomic slides from an ER+ breast cancer patient coloured by the activity scores of **(i)** luminal, **(ii)** epithelial, **(iii)** hallmark estrogen response, and **(iv)** hallmark E2F target genes. Red represents a higher activity score, and blue represents a lower activity score. The arrows point to areas of specific interest based on differences in heterogeneity of the different biological pathways. **C)** Spatial transcriptomic slides from an TNBC breast cancer patient coloured by the activity scores of **(i)** luminal, **(ii)** epithelial, **(iii)** hallmark estrogen response, **(iv)** basal, **(v)** mesenchymal, and **(vi)** pEMT gene sets. Red represents a higher activity score, and blue represents a lower activity score. The arrows point to areas of specific interest based on differences in heterogeneity of the different biological pathways.

We observed spatial heterogeneity in epithelial and luminal scores in ER+ tumors (**Fig 4B, i-ii**) but the signatures relating to basal, mesenchymal, and pEMT phenotypes were largely absent (**Fig S2D**). More specifically, we observed that even though different areas within the tumor displayed a similar extent of luminal signature they had varied epithelial scores (arrows shown in **Fig 4B, i-ii**). This observation is particularly important in the context of our survival analysis highlighting that EPI+LUM- or EPI-LUM+ phenotype show significantly worse survival compared to EPI+ LUM+ (**Fig 1D, ii**). We also investigated the spatial activity patterns of ER-driven genes that are generally found to be concordant with luminal and epithelial cell-states (Bouris et al., 2015; Lin et al., 2004). We observed that similar to luminal and epithelial scoring patterns, spatial transcriptomic slides had heterogeneous activity levels of ER response genes and E2F targets (a proxy for cell cycle activity) (**Fig 4B, iii-iv**). Such patterns of heterogeneity can dramatically impact sensitivity to anti-ER drugs (Chang, 2012). Thus, the degree of underlying heterogeneity of these molecular programs may limit the efficacy and evolution of resistance in ER+ breast cancer.

Next, we analysed spatial transcriptomics data from breast cancer patients classified to have basal disease. In one such patient sample, we noticed a more heterogenous expression of luminal and basal scores across the tissue samples as compared to a previously analysed case of luminal disease. We observed that the areas of normal breast tissue were high for luminal, epithelial and ER response gene set scores (**Fig 4C, i-iii,** top left tissue section). However, the tissue slices of the tumor were significantly more enriched in basal scores (**Fig 4C, iv**), with marked variability in terms of their epithelial and mesenchymal natures (**Fig 4C, ii, v**). This trend supports our results showing higher diversity in tumor epithelial-hybrid-mesenchymal states among basal tumors as compared to a luminal breast cancer case. Further, the concordance between the basal and the pEMT signature (**Fig 4C, compare iv with vi**) was higher compared to that between the basal and mesenchymal signature (**Fig 4C, compare iv with v**) with the pEMT signature being a proper subset of the spatial sections enriched for the basal signature of the tissue sections. This may explain the earlier observed bulk expression patterns where pEMT signatures correlated more closely with the basal signature compared to a mesenchymal signature.

### Basal breast cancer cell lines and tumor samples have higher phenotypic heterogeneity in terms of EMT

Next, we focused on pinpointing molecular underpinnings that may explain the association of the basal signature with the pEMT program. This association can be explained primarily by two scenarios: a) the basal subtype is primarily composed of cells that are pEMT, or b) the basal subtype is comprised of separate populations of cells that are epithelial and mesenchymal, resulting in a higher pEMT signature.

To gain a more comprehensive understanding of subtype-specific heterogeneity with respect to EMT status, we analysed single-cell RNA-seq of 32 breast cancer cell lines spanning all the clinical subtypes (GSE173634) (Gambardella et al., 2022). A previous study (Grasset et al., 2022) showed that CDH1+VIM+ TNBC cells were enriched during invasion and had higher colony forming ability. Thus, we classified this single-cell data based on gene expression values of CDH1 and VIM. The distribution of the difference between the imputed expression values of VIM and CDH1 showed multiple peaks (**Fig 5A, i**). The extreme peaks were labelled as epithelial and mesenchymal while the intermediate ones were labelled as hybrid phenotypes. We observed that all cells belonging to luminal A and luminal B cell lines were epithelial in nature. However, the basal cell lines had varied heterogeneity patterns along EMT axis. The TNBC type A cells were found to be either epithelial or hybrid E/M in nature, while the TNBC type B ones belonged to epithelial, hybrid E/M and mesenchymal states (**Fig 5A, ii-iii**). Interestingly, the HER2 subtype of breast cancer was largely epithelial in nature but the basal-like cell line MCF12A was predominantly in a hybrid E/M state (**Fig 5A, ii-iii**). These results indicate that while the luminal subtype is constituted primarily from an epithelial phenotype, the basal subtypes are more heterogeneous in terms of their E/M status and more likely to harbour a more pEMT cell state.

**Figure 5:**
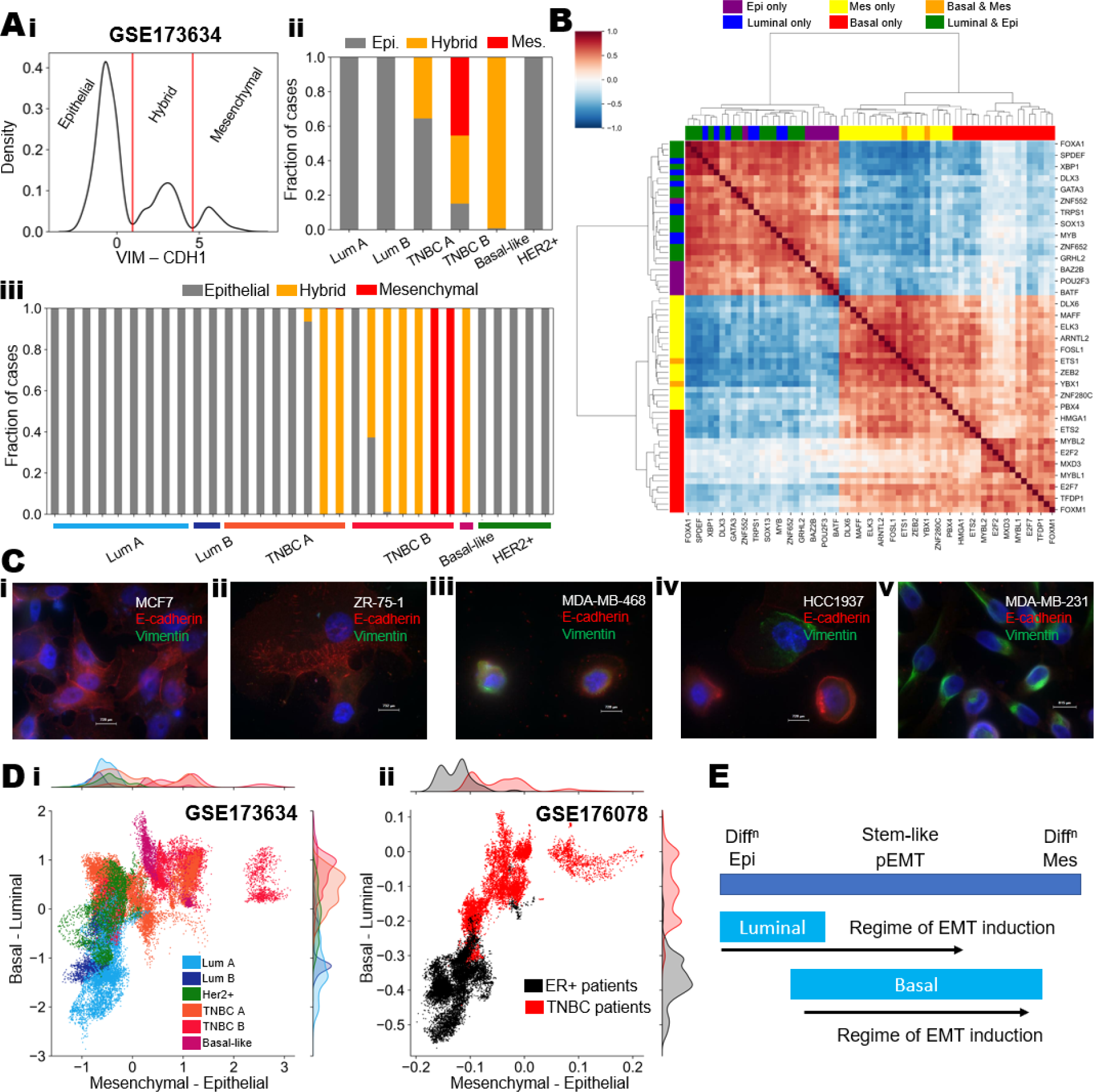
Heterogeneity patterns in single-cell analysis of breast cancer cell lines & patient derived cells. **A) i)** Kernel density estimate of VIM – CDH1 levels for cells belonging to all breast cancer cell lines (GSE173634). Red lines split the distribution into epithelial, mesenchymal and intermediate/hybrid states, based on difference in VIM and CDH1 levels. **ii)** Composition of different subtypes of breast cancer cell lines in terms of classified epithelial, mesenchymal and hybrid states. **iii)** Composition of different cell lines belonging to subtypes of breast cancer cell lines in terms of classified epithelial, mesenchymal and hybrid states. **B)** Gene-gene pairwise correlation heatmap showing top 20 transcription factors correlated with the luminal, basal, epithelial and mesenchymal signatures each in GSE173634. **C)** Immunofluorescence imaging of representative breast cancer cell lines showing E-cadherin and Vimentin levels. **D) i)** Scatterplot showing the distribution of single-cell RNA-seq data of breast cancer cell lines on the two-dimensional epithelial-mesenchymal status and luminal-basal status (GSE173634). x axis represents the epithelial-mesenchymal score (mesenchymal score – epithelial score) and y axis represents the luminal-basal score (basal score – luminal score). **ii)** Scatterplot showing the distribution of breast cancer cells from ER+ and TNBC patient samples on the two dimensional epithelial-mesenchymal status and luminal-basal status (GSE176078). x axis represents the epithelial-mesenchymal score (mesenchymal score – epithelial score); y axis represents the luminal-basal score (basal score – luminal score). **E)** Schematic showing the extent of heterogeneity and mapping of epithelial to mesenchymal transition status of luminal and basal breast cancer cases.

We analysed the top 20 transcription factors (TFs) that correlated with each of the four signatures luminal, basal, epithelial and mesenchymal (in GSE173634). Among the 20 TFs correlating with luminal and epithelial, 13 TFs were common, but among the 20 TFs correlating with mesenchymal and basal, only two TFs were common. These analyses suggest that the transcriptional programs that regulate luminal and epithelial states overlap while basal and mesenchymal states are regulated by distinct TFs. A hierarchically clustered pairwise correlation map of the top TFs, revealed antagonism between the luminal-epithelial and basal-mesenchymal group of TFs (**Fig 5B**). Furthermore, the basal and the mesenchymal transcription factors clustered separately with each other, indicative of the weak coupling between these two biological axes in contrast to the stronger associations of the luminal and epithelial TFs (**Fig 5B**).

Next, we probed whether these trends observed in RNA-seq data analysis could be recapitulated experimentally as well. Thus, we took 5 representative cell lines – 2 belonging to luminal subtype (MCF7 and ZR-75-1) and 3 belonging to basal subtype (HCC1937, MDA-MB-468 and MDA-MB-231). Each of these cell lines were probed for E-cadherin and Vimentin levels though immune-fluorescence experiments. It was observed that the luminal cell lines of MCF7 and ZR-75-1 were exclusively high for E-cadherin while have very little to no expression of Vimentin (**Fig 5C, i-ii**), indicating that the luminal cell lines were largely epithelial in nature. On the other hand, basal cell lines such as MDA-MB-468 and HCC1937 exhibited a mix of epithelial (high specifically for E-cadherin) and hybrid E/M phenotypes (co expression of E-cadherin and vimentin in the same cells) (**Fig 5C, iii-iv**). Finally, MDA-MB-231 showed largely mesenchymal cells (low E-cadherin, high Vimentin) with a few cells co-expressing E-cadherin and Vimentin (**Fig 5C, v**). The experimental results thus support our observations from bulk, single-cell and spatial transcriptomic data.

We next projected single-cell RNA-seq from both breast cancer cell lines and primary tumors on a two-dimensional plane, where the x-axis indicates EMT status (defined as difference in ssGSEA scores of mesenchymal and epithelial gene sets) and the y-axis denotes a difference between the ssGSEA scores of luminal and basal gene sets. For the 32 breast cancer cell lines (GSE173634), the trend between luminal-basal transition and EMT was non-linear; while a majority of the samples with high luminal scores clustered close to the epithelial end of the EMT axis, the basal-high samples spanned the entire EMT (**Fig 5D, i**). Similar patterns were observed from projection of single-cell RNA-seq data from tumor cells isolated from ER+ and TNBC patients (GSE 176078) (Wu et al., 2021) into this two-dimensional space. (**Fig 5D, ii**). These results collectively indicate that luminal cell lines (and ER+ tumors) were more restricted or homogenous in terms of their EMT state and exhibited predominantly an epithelial state while the basal breast cancer cell lines (and TNBC tumors) were not only more likely to exhibit a pEMT phenotype but also more heterogenous in terms of their EMT status (**Fig 5E**).

The enrichment of hybrid E/M phenotypes and/or higher heterogeneity along the EMT axis has been associated with worse survival in many cancer types (Jolly et al., 2022; Puram et al., 2017). Currently, no specific therapy targets either a hybrid E/M state or higher phenotypic heterogeneity, thus the enrichment of these attributes may explain the underlying mechanistic basis for difficulty in targeting basal-like tumors and TNBC.

### Mathematical modelling of gene regulatory networks captures phenotypic heterogeneity in breast cancer and pinpoints determinants of luminal-basal plasticity

Having uncovered the complex associations between the lineage characteristics of breast cancer cells and the status of cells along the epithelial-mesenchymal spectrum, we sought to understand the mechanistic underpinnings of these interconnected axes of plasticity. Specifically, we asked how breast cancer cells undergoing EMT may drive lineage plasticity and *vice versa*. Thus, we first assembled a gene regulatory network (GRN) based on experimental evidence (Table S4) to investigate the associations between the luminal-basal and epithelial-mesenchymal axes of cellular plasticity. This GRN is not inferred via statistical tools. It is expected to capture key factors involved in EMT and luminal-basal plasticity and to be capable of recapitulating underlying phenotypic heterogeneity and observed associations between different biological axes.

This GRN consists of representative TFs and genes associated with luminal (ERα66, PGR, GATA3 and FOXA1) (Bernardo et al., 2012, 2010; Eeckhoute et al., 2007; Kouros-Mehr et al., 2006), basal (SLUG and ΔNP63) (Nekulova et al., 2016; Storci et al., 2008), epithelial (CDH1 and miR-200) and mesenchymal (ZEB1, SLUG) phenotypes (Subbalakshmi et al., 2022), and regulatory interactions among them. We also incorporated two additional players: a) ERα36 as a marker for anti-ER therapy resistance (Thiebaut et al., 2020) and b) NRF2, as a reported stabilizer of the hybrid E/M state and also associated with drug resistance though its impact on cellular metabolism (Bocci et al., 2019; Kim et al., 2008; Wang et al., 2008) (**Fig 6A**). To understand the emergent dynamics of this GRN, we used RACIPE (Huang et al., 2017), a computational framework to identify the possible phenotypic space for a given GRN. RACIPE uses a set of coupled ordinary differential equations to simulate the dynamics of interconnected nodes (via regulatory links/edges) in a GRN and outputs the different possible steady state values of all nodes in the network. The relative expression levels of different genes/nodes constitute different cell-states that can correspond to observed phenotypes in breast cancer cell state heterogeneity.

**Figure 6:**
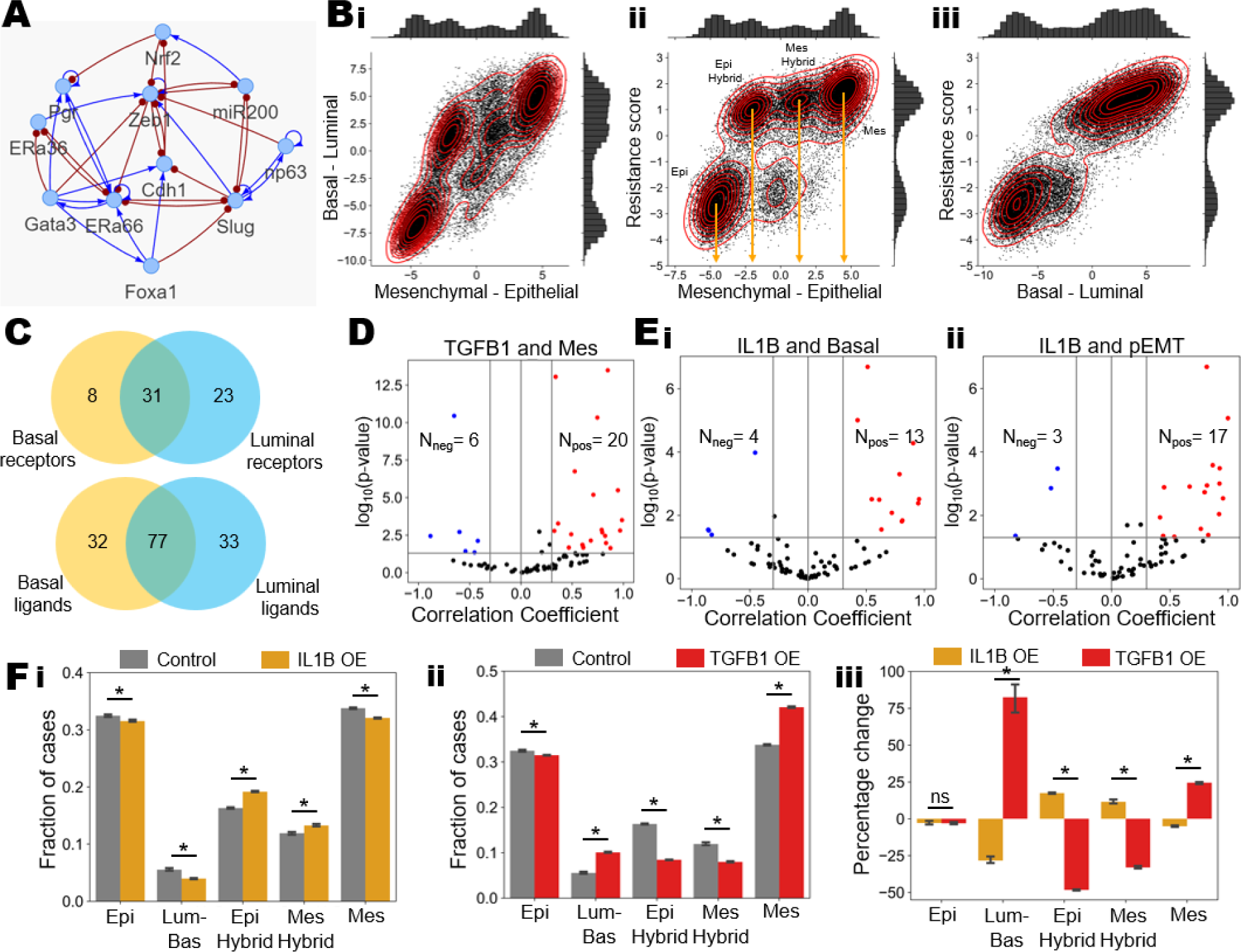
Gene regulatory network analysis for coupled luminal-basal and epithelial-mesenchymal plasticity. **A)** Gene regulatory network showing the representative genes from luminal, basal, epithelial and mesenchymal programs and the associated regulatory links between them. Blue links represent activation, while red links represent inhibition. **B)** Simulated steady state solutions projected on a two dimensional **i)** epithelial-mesenchymal axis and luminal-basal axis, **ii)** epithelial-mesenchymal axis and resistance score, and **iii)** luminal-basal axis and the resistance score. **C)** Number of common and unique receptors that were associated with luminal and basal cell type and respective tumor microenvironments. **D)** Meta analysis of 80 breast cancer specific datasets showing volcano plots of correlations for the TGFB1 gene and the mesenchymal program. Each dot denotes a unique GSE ID (dataset). Pearson’s correlation coefficients and the corresponding p-values are mentioned. **E)** Meta analysis of 80 breast cancer specific datasets showing volcano plots of correlations for IL1B gene expression with **i)** basal signature and **ii)** the pEMT program. Pearson’s correlation coefficients and corresponding p-values are indicated. **F)** Simulation results showing the fraction of steady state solutions belonging to the different cell states in control and **i)** IL1B over-expression (OE) and **ii)** TGFB1 OE scenarios. **iii)** Simulation results showing percentage change in each observed phenotype compared to the control scenario, in TGFB1 and IL1B OE scenarios.

We defined the following scores to better understand the association between EMT and luminal-basal plasticity: a) luminal score as the sum of normalised steady state values of ERα66, GATA3, PGR, and FOXA1, b) basal score as the sum of normalised steady state values of ΔNP63 and SLUG, c) epithelial score as the sum of normalised steady state values of CDH1 and miR-200, d) mesenchymal score as the sum of normalised steady state values of ZEB1 and SLUG, and e) resistance score as the difference between normalised steady state levels of ERα36 and ERα66. We observed that the epithelial-mesenchymal score (= mesenchymal score – epithelial score) was multimodal in nature with two hybrid states (referred to as epithelial-hybrid and mesenchymal-hybrid) in addition to the canonical epithelial and mesenchymal states. The luminal-basal score (= basal score – luminal score) distribution was largely trimodal (**Fig S2E**). Thus, our GRN dynamics can recapitulate the multiple hybrid E/M phenotypes that have been well-reported experimentally (Brown et al., 2022; Jolly et al., 2022), as well as a lumino-basal phenotype identified recently in breast cancer (Fan et al., 2020; Mohamed et al., 2023).

Upon projecting these simulated node values on a two-dimensional plane of luminal-basal and epithelial-mesenchymal axes, we could recapitulate the non-linear relationship between the epithelial-mesenchymal axis with the luminal-basal axis (**Fig 6B, i**) as observed for breast cancer cell lines (**Fig 5D, i**) and patient-derived tumor cell data (**Fig 5D, ii**) at a qualitative level. More specifically, we observed that while the solutions corresponding to a luminal state are almost exclusively epithelial in nature, those that correspond to a basal like state could be epithelial, mesenchymal or hybrid E/M (pEMT) on the epithelial-mesenchymal spectrum (**Fig 6B, i**). This observation suggests that the GRN considered here can explain and reproduce the major associations noted in extensive multi-modal transcriptomic data (bulk, single-cell, spatial) analysis across breast cancer cell lines and tumors.

Further, our simulation results highlight that the most epithelial cluster is predominantly low in resistance score, i.e. being sensitive to anti-ER drugs such as tamoxifen. However, the other three clusters (epithelial-hybrid, mesenchymal-hybrid and mesenchymal) had significantly higher resistance scores (**Fig 6B, ii**). These model predictions are reminiscent of prior experimental observations that partial and/or full EMT can drive resistance to tamoxifen and *vice versa* (Kastrati et al., 2020; Sahoo et al., 2021a). We also observed that the luminal-basal status strongly correlated with the resistance score, suggesting that lineage determination is a crucial factor for sensitivity to anti-ER drugs (**Fig 6B, iii**). We found that the two hybrid E/M clusters had higher levels of NRF2 (**Fig S2F**), consistent with the reported literature (Bocci et al., 2019; Kim et al., 2018; Pasani et al., 2020). These results indicate that the GRN considered here can sufficiently capture the observed associative trends in breast cancer and further correlate luminal-basal and epithelial-mesenchymal trends to their corresponding sensitivity to anti-estrogen drugs such as tamoxifen.

Having simulated a GRN that couples the biological axes of lineage characteristics (luminal/basal), epithelial/mesenchymal status and the degree of sensitivity to the widely used anti-ER drugs, we wished to identify potential ligands and receptors that could drive a luminal to basal phenotypic switch, thus limiting the efficacy of the anti-ER drugs especially in the context of ER+ breast cancer. To do this we analysed a single-cell patient atlas of 7 ER+ breast cancer patients and 5 TNBC patients (GSE176078). Each of these samples had annotated tumor cells as well as corresponding stromal and immune cell types. Thus, we used LIANA pipeline (Dimitrov et al., 2022) to score for top ligands and receptors for each patient separately. Specifically, we identified the ligands and receptors that were specific to the target cell type pre-annotated to be luminal A or basal subtype of cancer cells in ER+ and TNBC patients, respectively. We focused on ligands and receptors that were consistently expressed in all patient samples with at least 100 target breast cancer cells. We created separate lists for ligands and receptors for luminal A and for basal breast cancer cells; for instance, the top receptors unique to luminal A tumor cells contained ESR1, MUC1 and ERBB3, each of which has been associated with a luminal subtype (Balko et al., 2012; Semba et al., 2023). We observed substantial overlap between the ligand and receptors from the basal and luminal cell types (**Fig 6C**), suggesting potential overlaps in signalling between luminal and basal cell types with the tumor microenvironment.

Next, we focused on the list of common ligands as they can potentially act on luminal cells and cause a transition to a basal-like phenotype. TGF-β1 was one such common ligand, which is a well-known driver of EMT in many carcinomas. To assess how likely a ligand is to affect the luminal-basal phenotype of the cells, we performed meta-analysis of each of the common ligands in 80 bulk transcriptomics datasets from breast cancer. We found that TGFβ1 was amongst the top genes that correlated positively with a more mesenchymal phenotype (**Fig 6D**). On the other hand, we found IL-1β to be one of the top genes to be correlated with a basal phenotype (**Fig 6E, i**) as well as pEMT phenotype (**Fig 6E, ii**). Further, we interrogated whether our GRN could reproduce the plasticity patterns that can be driven by TGF-β1 or IL-1 β. To do this we extended our GRN (**Table S4**) to include IL-1β as well as TGFβ and performed simulations to over express these nodes *in silico*. Our model predicted that IL-1β overexpression led to a significant increase in the population of the epithelial-hybrid and the mesenchymal-hybrid populations, with a concomitant decrease in the frequency of epithelial and mesenchymal phenotypes (**Fig 6F, i, iii**). On the other hand, TGF-β1 over expression caused a marked increase in the frequency of mesenchymal phenotype, and a simultaneous reduction in epithelial, epithelial-hybrid and mesenchymal-hybrid states (**Fig 6F, ii-iii**). There was also a significant increase in the proportion of the luminal basal hybrid phenotype (**Fig 6F, ii-iii**). Although both IL-1β and TGF-β1 cause a net decrease in the epithelial phenotype and increase the mesenchymal nature, their impact is quite distinct: IL-1β can enrich the hybrid E/M phenotypes that are basal in nature. This prediction is consistent with recent experimental observations about the impact of IL-1β *in vitro* and *in vivo*: a) IL-1β treatment can induce two stabilizers of hybrid E/M phenotype NRF2 and SLUG (González-Carnicero et al., 2023; Li et al., 2020), b) IL-1β treatment of MCF7 luminal breast cancer cells can induce ΔNP63 and mediate subsequent therapy resistance (Mendoza-Rodríguez et al., 2019), and c) IL-1β treatment can prevent differentiation of metastatic-initiating cells to highly proliferative epithelial cells, inhibiting overt metastatic growth (Castaño et al., 2018). Together, our analysis suggests IL-1β to be a potent target to prevent luminal-to-basal lineage plasticity.

Overall, our simulation results show that the core GRN modelled here can capture complex associations between the luminal-basal and epithelial-mesenchymal axes and explain underlying mechanisms of the phenotypic heterogeneity in basal breast cancer. This GRN can also serve as a robust framework to simulate the effect of other signalling molecules to enrich for different phenotypes as a consequence of emergent properties of cross-linked feedback loops among different factors.

## Discussion

Phenotypic heterogeneity is a fundamental feature of biological systems implicated in better chances of survival of a population under various dynamically varying environmental stress levels (van Boxtel et al., 2017). Not surprisingly, cancer cells are heterogenous along various functional and molecular axes. This heterogeneity often helps them to evade therapeutic attacks and adapt to their changing environments, eventually driving their metastatic dissemination and colonization (Brown et al., 2022; Chauhan et al., 2021; Pastushenko et al., 2018; Rambow et al., 2019; Yu et al., 2013). Single-cell approaches have been instrumental in characterizing such heterogeneity, but usually along one axis (Deshmukh et al., 2021; Jia et al., 2019; Karacosta et al., 2019). Analysing interconnections among different axes of heterogeneity is relatively poorly understood.

Here, we used a multi-modal (single-cell, bulk and spatial) transcriptomic data analysis approach to identify the associations between two key axes of heterogeneity in breast cancer – EMT and luminal-basal (lineage) plasticity (Beltran et al., 2019; Kern et al., 2022; Mohamed et al., 2023; Padua et al., 2018). Often, these axes have been assumed to be synonymous or largely overlapping, but many questions remained unanswered: a) how do luminal and basal features map to epithelial, hybrid E/M and mesenchymal phenotypes?; b) which breast cancer subtype (luminal/basal) have higher heterogeneity in terms of EMT?; and c) what feedback loops connect these two plasticity axes?

These analyses of breast cancer cell lines and primary tumors have unravelled that while luminal breast cancer samples are predominantly epithelial and quite homogenous, basal breast cancer samples correspond to hybrid E/M state(s) with more phenotypic heterogeneity along the EMT spectrum. Both of these hallmark features of basal breast cancer have independently been associated with worse patient survival in many cancer types (Brown et al., 2022; Jolly et al., 2022; Vipparthi et al., 2022). Thus, their co-existence can possibly explain the aggressive behaviour and limited therapeutic response of basal-like breast cancers (Milioli et al., 2017). Further, given the relatively higher plasticity of hybrid E/M phenotypes as compared to ‘fully epithelial’ or ‘fully mesenchymal’ phenotypes (Hari et al., 2022; Kröger et al., 2019), the presence of hybrid E/M phenotypes can facilitate more phenotypic heterogeneity in a population. Our results are consistent with observations of higher PD-L1 protein levels (Ali et al., 2015) and tamoxifen resistance (Fan et al., 2020) in basal-like tumors, given that both these traits have been previously linked to hybrid E/M cells (Sahoo et al., 2021a, 2021b). Even among luminal tumors, those expressing low levels of ER can display higher basal-like phenotypes (Mohamed et al., 2023), endorsing previous results that *ESR1* expression closely associates with luminal breast cancer cells and that silencing ER can drive EMT (Al Saleh et al., 2011). Because EMT and tamoxifen resistance can both drive each other (Hiscox et al., 2006; Sahoo et al., 2021a), our results suggest that tamoxifen resistance can govern lineage plasticity, i.e., luminal-to-basal switch, as well. Similar interconnections about lineage plasticity, EMT and anti-androgen resistance have been reported in prostate cancer (Jindal et al., 2023; Mu et al., 2017) and small cell lung cancer (Groves et al., 2023). These correspondences indicate possible generalisability of our results to other cancer types as well.

Lineage plasticity is being increasingly reported in the context of breast cancer, but with limited mechanistic understanding. For instance, mature mammary luminal epithelial cells can give rise to Krt14 and Sox9-expressing basal-like carcinomas that can metastasize (Kern et al., 2022). Similarly, basal-like tumorigenesis involves luminal-to-basal reprogramming with gain in stemness (Christin et al., 2020; Landragin et al., 2022). Further, luminal-basal hybrid cells can express NP63 (basal marker) while maintaining functional levels of ER-alpha (associated with luminal phenotype) (Padua et al., 2018). Our systems-level analysis integrating multi-modal transcriptomic data with mechanism-based models for underlying regulatory networks explains these *in vitro* and *in vivo* observations, reveals hallmarks of basal breast cancer in terms of EMT, and offers a predictive platform to better characterize and control intra-tumor phenotypic heterogeneity in breast cancer.

## METHODS

### ssGSEA scores for bulk transcriptomics

We used previously published signatures for luminal and basal breast cancer (Nair et al., 2022), for epithelial and mesenchymal state (Tan et al., 2014) and for pEMT state (Puram et al., 2017) Hallmark signatures (estrogen response, E2F target genes) were taken from MSigDB database (Liberzon et al., 2011). ssGSEA scores were calculated for bulk transcriptomic samples using the gseapy python package (Fang et al., 2023) to estimate the activity of biological pathway of interest. A correlation was considered significant if the Pearson’s correlation coefficient is greater than 0.3 or lesser than 0.3 with a p-value lesser than 0.05. Meta analysis was performed on a list of 80 breast cancer specific bulk RNA/microarray transcriptomic datasets (**Table S3**).

### Survival analysis

Overall survival data was acquired from TCGA. Based on the median of sample scores, all samples were split into 4 groups: epithelial-high mesenchymal-low (EPI+MES-) (reference group), epithelial-high mesenchymal-high (EPI+MES+), epithelial-low mesenchymal-high (EPI-MES+), and epithelial-low mesenchymal-low (EPI-MES-). Similarly, epithelial-high luminal-high (EPI+LUM+) (reference group), epithelial-high luminal-low (EPI+LUM-), epithelial-low luminal-high (EPI-LUM+), and epithelial-low luminal-low (EPI-LUM-). The R package ‘survival’ was employed to perform the Kaplan–Meier analysis. Reported p-values were calculated using a log-rank test. Cox regression was used to determine the hazard ratio (HR) and confidence interval (95% CI) for TCGA cohorts, and forest plots were made using ‘ggforest’ function from ‘survminer’ package.

### Methylation data Analysis

Methylation data (beta values) from GSE42944 for breast cancer cell lines was considered. The beta values for CpG islands vary between 0 (unmethylated) and 1 (fully methylated). The heatmap included only previously identified epithelial and mesenchymal genes (Tan et al., 2014) and cell lines included in CCLE breast cancer cohort to facilitate direct comparison between the RNA-Seq and methylation data. We further performed z-normalisation and scaled the values between 0 and 1 to portray only the relative amount of methylation values for each gene across the cell lines.

### Spatial Transcriptomics data analysis

Spatial transcriptomics datasets in the public domain for 6 patients (2 estrogen receptor positive and 4 triple negative patients) (Wu et al., 2021) were reanalysed for assessing the activity of various biological pathways. Count matrices were first imputed by MAGIC algorithm (van Dijk et al., 2018) and activity scores were calculated on imputed values using AUCell (Aibar et al., 2017). Pre-processing of spatial data and images was done as per the Seurat pipeline (Hao et al., 2021).

### Cell Lines and culture

Cell lines MDA-MB-231, MCF7 and HCC1937 were obtained from the American Type Culture Collection (ATCC-Manassas, VA). MDA-MB-468 and ZR-75-1 were obtained from NCCS (Pune, India) where cell authentication was performed using STR profiling. MDA-MB-231 and MDA-MB-468 were maintained in L-15 (Leibovitz) medium (Sigma-Aldrich), MCF7 in DMEM-Hi Glucose medium (Sigma-Aldrich) HCC1937 and ZR-75-1 in RPMI 1640 media (Gibco), HEPES buffered and supplemented with 10% (v/v) heat inactivated Fetal Bovine Serum (Himedia) and 100 U/ml penicillin and streptomycin (Gibco). All cells were maintained in a humidified incubator with 5% CO2 at 37°C except for MDA-MB-231 and MDA-MB-468 that were maintained with 0% CO2. For all experimental assays using cell lines, a passage number below 20 was used and all cell lines were subjected to frequent recharacterization by immunophenotyping and testing of mycoplasma.

### Dual immunofluorescence

Cells (1x10^4^) were seeded on poly-L-lysine coated coverslips. Immunofluorescence was performed as reported previously (Nair et al., 2016) by incubating cells in primary antibodies-anti-E-cadherin (Abcam-EP700Y) and anti-Vimentin (BioGenex) overnight at 4°C at specific dilutions-1:500 and 1:25 respectively. This was followed by labelling with specific secondary antibodies - Alexa Fluor® 488 Chicken Anti-Mouse IgG (H+L) for Anti-Vimentin and Alexa Fluor 568 Donkey Anti-Rabbit IgG for anti-E-cadherin for 1 h at room temperature. The slide was then mounted on gold antifade reagent with DAPI and examined under a fluorescent microscope (Olympus BX51).

### Single cell RNA sequencing and cell-cell communication data analysis

Count matrices for single-cell RNA sequencing data were imputed by MAGIC algorithm (van Dijk et al., 2018). Activity scores were calculated on imputed values using AUCell (Aibar et al., 2017). Lists of top cell-cell communication receptor ligand pairs were estimated using the LIANA package (Dimitrov et al., 2022). For estimating the top ligand receptor pairs, pre-labelled luminal A and basal cell types from each patient were considered and all other cell types including themselves were considered as potential ligand producing cells while the receptors were assumed to be only expressed on the chosen luminal A or basal subtype of cells. Only those ligand receptor pairs which had a cellphonedb p-value < 0.05 and sca LRscore > 0.8 and were expressed in all ER+ or TNBC were considered as unique/common ligands/receptors. Ligands or receptors that were only specific to luminal cells in ER+ breast cancer patients were considered to be luminal specific ligands/ receptors. Similar analysis was done for basal cells in TNBC as well. Top genes were subjected to meta-analysis with relevant pathways & their expression in bulk transcriptomics.

### RACIPE simulations

Random Circuit Perturbation (RACIPE) was employed to generate an ensemble of kinetic models for a given GRN. The GRN contains nodes and edges (inhibitory or activating) among them. The dynamics of each node was determined using a set of coupled ordinary differential equations (ODEs) (Huang et al., 2017). Each node/gene had basal production and degradation rates as ODE parameters. Shifted Hill functions were multiplied to the production rate to incorporate the effects of excitatory and inhibitory links incoming to that node/gene. All steady-state values obtained from RACIPE, which were initially in log2 scale, were converted into z-scores, to indicate relative levels. RACIPE simulations were done in triplicates, each replicate with 10,000 parameter sets, and 100 initial conditions for each parameter set. Euler’s Method was employed for numerical integration. RACIPE chooses kinetic parameters from a large range of biologically realistic parameter values to identify a majority of states that are allowed by a given GRN. A single RACIPE parameter set and associated random initial conditions has the potential to produce one or more stable steady-state solutions. However, for this analysis, up to six stable steady-state solutions were considered.

The luminal score was calculated as sum of normalised steady state values of ERα66, GATA3, PGR, and FOXA1. Similarly, basal score included ΔNP63 and SLUG; epithelial score incorporated CDH1 and miR-200 and the mesenchymal score consisted of ZEB1 and SLUG. Furthermore, the resistance (to anti-ER drugs) score was calculated as the difference between steady-state values of ERα36 and ERα66. Additionally, RACIPE was used to perform overexpression (OE; 100x) of TGFB1 & IL1B genes separately for a modified GRN consisting of both these genes and the steady state results obtained were then compared with control (no OE) RACIPE simulations.

### MINT-Chip data analysis

MINT-Chip data for 4 breast cancer cell lines (MCF7, ZR-75-1, HCC38 and HMLER) were processed to obtain the enriched promoters (considered to be 5000 base pairs either side of the transcriptional start site) for different cell lines and CD44 status. First, a list of top correlated epithelial and mesenchymal genes (Spearman’s correlation coefficient > 0.5 and p-value < 0.05) were obtained from RNA seq data of the same cell lines (GSE184647). Amongst these genes which were labelled to be epithelial or mesenchymal, the proportion of genes that had an enriched promoter in either activation (H3K27ac) or inhibitory (H3K27me3) marks were quantified and compared across the cell lines or cell lines with specific CD44 status.

## Conflict of Interest

The authors declare no conflicts of interest.

## Author contributions

Conceptualized and designed research: MKJ

Supervised research: CLC, JSP, JAS, MKJ

Performed research: SS, SR, MGN, MP, BPSN, SM, CMN, ADM, HS, AGN

Interpreted data: SS, SR, MGN, MP, BPSN, SM, CMN, ADM, CLC, JSP, JAS, MKJ

Funding acquisition: JSP, JAS, MKJ

Manuscript writing/editing: SS (prepared first draft), JAS, MKJ (edited with inputs from all authors)

## Funding

This work was supported by Ramanujan Fellowship awarded by SERB (Science and Engineering Research Board), Department of Science and Technology (DST), Government of India, awarded to MKJ (SB/S2/RJN-049/2018). SS is supported by PMRF (Prime Ministers Research Fellowship) awarded by DST, Government of India. JASs is supported by NCI 1R01CA233585-04.

## Supplementary Figures

**Figure S1:**
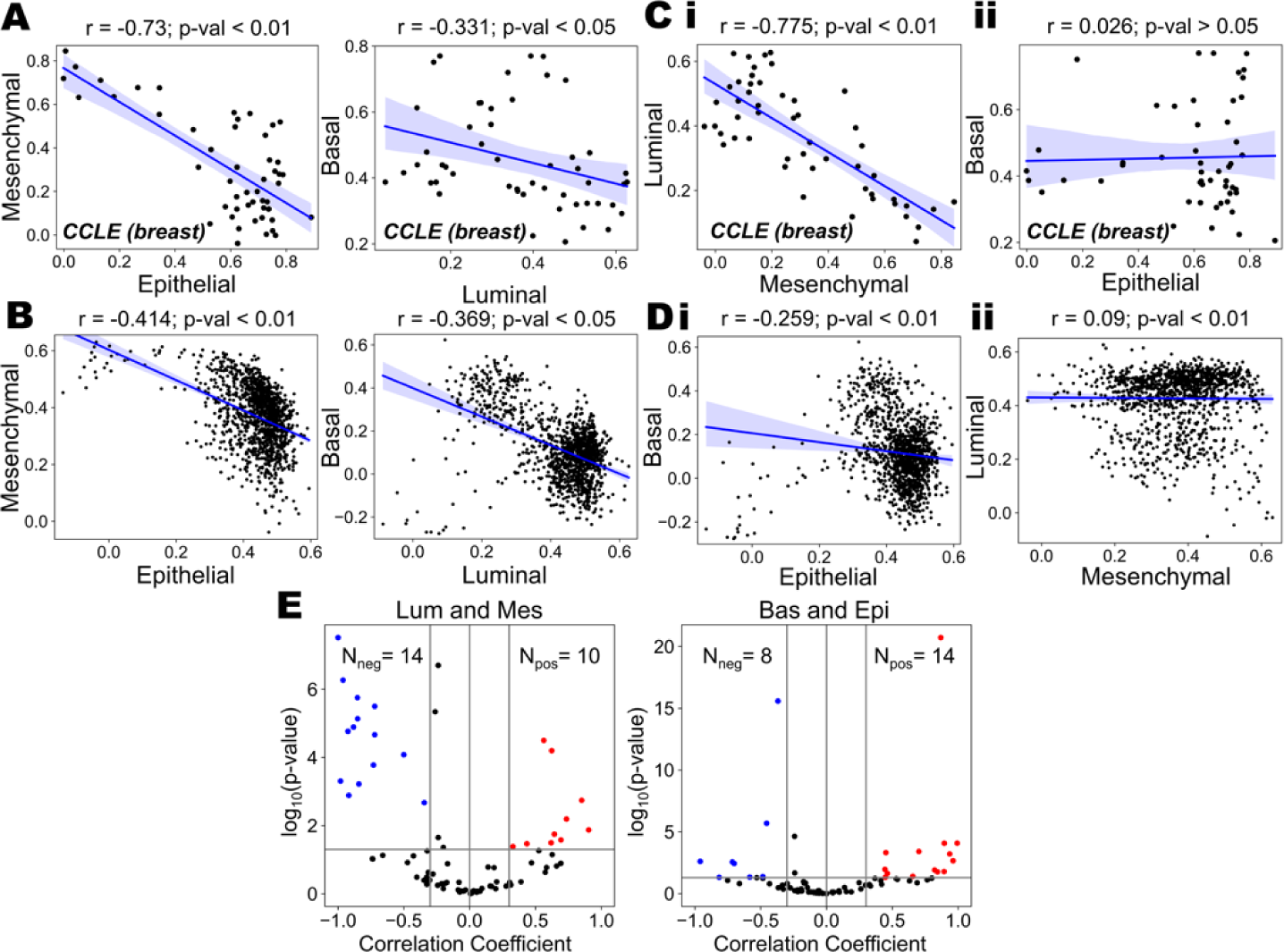
Associations between the luminal-mesenchymal and basal-epithelial programs in breast cancer. **A)** Scatterplots showing the correlations between the ssGSEA scores of i) Epithelial-Mesenchymal and ii) Luminal-Basal programs in CCLE breast cancer cell lines. Pearson’s correlation coefficients have been reported with corresponding p-values. **B)** Scatterplots showing the correlations between the ssGSEA scores of i) Epithelial-Mesenchymal and ii) Luminal-Basal programs in TCGA breast cancer patients. Pearson’s correlation coefficients have been reported with corresponding p-values. **C)** Scatterplots showing the correlations between the ssGSEA scores of i) Mesenchymal-Luminal and ii) Basal-Epithelial programs in CCLE breast cancer cell lines. Pearson’s correlation coefficients have been reported with corresponding p-values. **D)** Scatterplots showing the correlations between ssGSEA scores of i) Mesenchymal-Luminal and ii) Basal-Epithelial programs in TCGA breast cancer patients. Pearson’s correlation coefficients have been reported with corresponding p-values. **E)** Meta analysis of 80 breast cancer specific datasets showing volcano plots of correlations for i) Mesenchymal-Luminal and ii) Basal-Epithelial programs. Pearson’s correlation coefficients and corresponding p-values are given.

**Figure S2:**
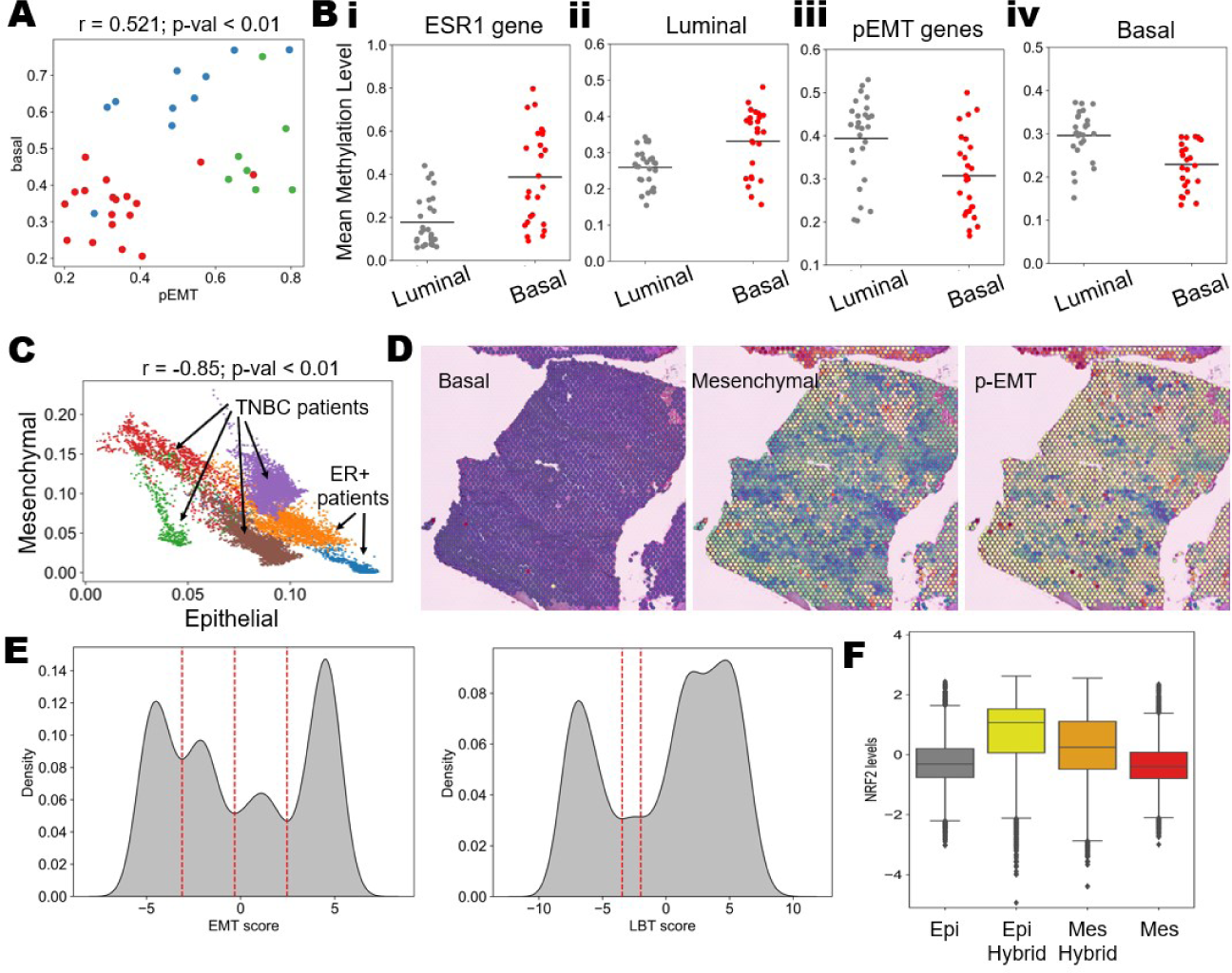
Epigenetic, Spatial and Simulation results of breast cancer heterogeneity. **A)** Scatterplots showing correlations between the ssGSEA scores of pEMT-basal programs in breast cancer cell lines. Pearson’s correlation coefficients have been reported with corresponding p-values. **B)** Comparison of mean methylation levels across luminal and basal breast cancer for i) ESR1 gene ii) Luminal genes iii) pEMT genes and iv) Basal genes. **C)** Scatterplot showing position of the ER+ and TNBC patients on a two dimensional epithelial-mesenchymal plane. Each point on the plane is based on gene expression values for a specific spot on the spatial transcriptomic datasets. **D)** Spatial transcriptomic slides from an ER+ breast cancer patient coloured by the activity scores of basal, mesenchymal and pEMT programs. Red represents a higher activity score, and blue represents a lower activity score. **E)** Kernel density estimate of epithelial-mesenchymal score and luminal-basal scores from RACIPE simulations. Red dotted lines show the position of minima used to define phenotypes. **F)** Simulated gene expression values for NRF2 gene in different EM phenotypes.

## Notes

### Competing Interest Statement

The authors have declared no competing interest.

